# Investigating neural speech processing with functional near infrared spectroscopy: considerations for temporal response functions

**DOI:** 10.64898/2026.03.20.713212

**Authors:** Johanna Wilroth, Nancy Sotero Silva, Ali Tafakkor, Bruno de Avo Mesquita, Emily Y. J. Ip, Bonnie Lau, Jaimy Hannah, Giovanni M. Di Liberto

**Author notes:** These authors contributed equally as senior authors (Jaimy Hannah, Giovanni M. Di Liberto).

## Abstract

Functional near infrared spectroscopy (fNIRS) is increasingly used in hearing and communication research, with advantages such as robustness to movement artifacts, improved spatial resolution, and flexibility of contexts in which it can be applied. At the same time, the field is progressively moving towards more continuous, naturalistic listening paradigms resulting in the widespread adoption of speech tracking analyses such as temporal response functions (TRFs) in electroencephalography (EEG) and magnetoencephalography (MEG) studies. However, it remains unclear whether these analyses can be applied to slower haemodynamic signals measured by fNIRS. In the present study, we investigated whether a TRF framework can similarly be applied to fNIRS data recorded during continuous speech perception. Eight participants listened to speech simultaneously while fNIRS signals were acquired in a hyperscanning setup. Speech features were regressed onto the haemodynamic responses to test the feasibility and interpretability of fNIRS-based TRFs. Prediction correlations between observed and modelled fNIRS signals across speech features were higher than those typically reported for EEG- and comparable to those reported for MEG-TRF studies. Moreover, these correlations did not overlap with a null distribution generated from triallJmismatched fNIRS data, confirming statistical significance and were slightly greater than those obtained from a conventional GLM approach. Our findings support that TRF estimation method can yield meaningful and statistically significant responses from fNIRS data.

**Highlights:**

- TRF modelling can be meaningfully applied to fNIRS data acquired during speech listening tasks.
- Prediction correlations between actual and modelled fNIRS signals were above chance level, with values comparable to previous EEG/MEG studies.
- TRFs explained more fNIRS variance than a conventional GLM approach.

## 1. Introduction

Neurophysiology research on speech perception has traditionally relied on highly controlled experimental paradigms that use short, isolated units of speech such as syllables, words, or brief sentences. While these approaches allow a high degree of control and precise alignment with event-related designs, they fail to capture important properties of natural speech, including continuity, temporal dynamics, and contextual complexity (Shen et al., 2024). More recently, however, there has been a shift towards using continuous, naturalistic stimuli that better approximate real-life listening situations, motivated by the opportunity to investigate how social factors interact with sound and language processing (Di Liberto & Ip, 2025; Hu et al., 2025; Rowland et al., 2018; Sonkusare et al., 2019). While traditional electrophysiological paradigms primarily rely on stimuli on the order of milliseconds (Rossi et al., 2012), several neuroimaging studies have demonstrated the feasibility of using longer speech segments (e.g., audio-books; podcasts) to study speech comprehension and listening effort (Bálint et al., 2025; Bertachini et al., 2021; Levin et al., 2022; Pollonini et al., 2014; van de Rijt et al., 2016).

This shift toward continuous speech has been accompanied by the introduction of analytic frameworks for modelling the relationship between continuous stimulus features and neural responses. A widely adopted framework for estimating such relationships is the Temporal Response Function (TRF), where encoding models are derived using regularised ridge regression (Crosse et al., 2016; Crosse et al., 2021). TRFs have been extensively applied to electroencephalography (EEG) and magnetoencephalography (MEG) data to quantify neural tracking of speech and music, shedding light on the brain represents linguistic and acoustic units (Brodbeck et al., 2018; Di Liberto et al., 2023; Menn et al., 2023), implements auditory attention mechanisms (Ding & Simon, 2012; O’Sullivan et al., 2015), integrates audiovisual cues (Crosse et al., 2015), and implements predictive processing (Di Liberto et al., 2018), among other cognitive operations (Di Liberto et al., 2020).

EEG and MEG offer excellent temporal resolution and remain indispensable for studying the millisecond-level dynamics of speech perception and language processing. However, like all neuroimaging techniques, they come with modality specific trade-offs that can make certain naturalistic paradigms more suitable than others. For instance, the low spatial resolution of EEG limits precise source localisation, especially for deeper or distributed cortical generators. MEG offers improved spatial resolution to EEG, but both modalities are sensitive to movement-related artifacts, which can constrain experimental designs involving interactive, mobile, or highly naturalistic behaviours. This is especially true with MEG, where magnetic shielding is critical, and so participation occurs in small shielded in rooms, with certain MEG systems also requiring participants to be head fixed. For these reasons, considering other neuroimaging methods with better experimental flexibility is important to broaden the range of paradigms that can be meaningfully investigated.

Functional near-infrared spectroscopy (fNIRS) is an optical neuroimaging technique that indirectly measures brain activity (Czeszumski et al., 2020; Eulau & Hirsh-Pasek, 2024; Quaresima et al., 2019). It relies on the comparison of haemoglobin oxygenation concentrations in nervous tissues, offering blood flow measurements (Eulau & Hirsh-Pasek, 2024; Pinti et al., 2020). Due to its non-invasive nature and robustness to movement artifacts, it has been used in substitution or combination with EEG, reaching cortical areas up to 1.5 to 2 cm in depth (Czeszumski et al., 2020; Eulau & Hirsh-Pasek, 2024; Pinti et al., 2020). Additionally, due to its compatibility with electronic and magnetic devices, fNIRS has also been widely used in research focused on hearing and speech perception, especially with hearing aid and cochlear implant users (Anderson et al., 2017; Bell et al., 2020; Sevy et al., 2010).

fNIRS has been successfully employed to study the speech processing hierarchy in infants and adults (Rossi et al., 2012), including phonemic contrasts to syllables and words (Minagawa-Kawai et al., 2008), sentence-level processing (Peña et al., 2003), and challenging listening conditions such as vocoded speech (Lawrence et al., 2018). However, these studies typically rely on comparing the condition of interest with a baseline condition using general linear models (GLM). This reliance stems from the slow and overlapping nature of the haemodynamic response, which peak several seconds after stimulus onset, making it difficult to isolate neural responses without clearly separated blocks. Such block designs place constraints on experimental flexibility, limiting the study of natural speech processing.

Here, we examine whether a TRF framework can be meaningfully applied to fNIRS data acquired during a continuous speech listening task. The fNIRS experiment was conducted as part of the 1^st^ Cognition and Natural Sensory Processing (CNSP) hackathon, a satellite event to the 8^th^ International Conference on Auditory Cortex. In a single hyperscanning session using a classroom-style design, fNIRS data were recorded simultaneously from eight participants as they listened to podcast dialogues presented through a loudspeaker. The stimuli were identical to those used in a previous EEG study on podcast dialogue listening (Ip et al., 2025), providing clear expectations for the temporal response patterns. Encoding TRF and decoding analyses were used to assess the extent to which fNIRS can capture speech-related neural processing across multiple levels of the processing hierarchy. Although this study was conducted in a hyperscanning context, this was not the primary focus of our analyses. Rather than targeting intersubject synchronisation, we aim to bridge TRF analyses with the increasing use of fNIRS in hearing and communication research, and to evaluate if fNIRS can support continuous, naturalistic paradigms using analysis methods designed for electrophysiology.

## 2. Methods

### 2.1 Participants

Eight volunteer Hackathon attendees (four female, four male) between 25 and 43 years of age (*M* = 30.63) took part in the experiment. Five of the eight participants were native English speakers and the remaining three were highly proficient in English. The study was approved by the School of Computer Science and Statistics Research Ethics Committee at the University of Dublin, Trinity College, and all participants provided written informed consent. Data collection took place in a workroom with natural lighting in the office building where Hackathon activities were carried out in Maastricht, the Netherlands.

### 2.2 Stimuli and Task

A previous experiment on non-simultaneous EEG recordings was adapted for the hyperscanning fNIRS setting (Ip et al., 2025), following a similar presentation of stimuli. Participants sat together in front of a screen where written instructions and a visual fixation cross were displayed. Audio stimuli were delivered by loudspeakers at the front of the room and presented at a sampling rate of 44,100 Hz. Stimuli were presented and controlled by the PsychoPy Python library version 2025.1.1 (Peirce et al., 2019). The audio samples consisted of 28 dialogues taken from four sources: two American podcasts (*Brains On!* and *Forever Ago*), a YouTube series by WIRED (*5 Levels of Difficulty*), and interviews with the hosts from the podcast *Forever Ago*. All podcasts and shows are publicly available online^1^. The dialogues discussed general interest topics related to science, arts, and childhood memories. All audios were one-on-one dialogues from either adult-adult (10 trials) or adult-child (18 trials) interactions, totalling approximately one hour. The whole experiment was run in a single session, as data from all participants were acquired simultaneously.

### 2.3 fNIRS Data Acquisition

fNIRS data was recorded with Brite Ultra System (Artinis Medical Systems B.V., Elst, The Netherlands), a system specifically designed for hyperscanning recordings. The fNIRS devices were wireless (model Brite MKII), each including 18 optodes (10 transmitters and 8 receivers) leading to 16 long-separation channels and two short-separation channels (see Table 1). Optodes were mounted in caps, covering frontal and temporal regions. The montage of the optodes and placement of the caps was carried out by the volunteers, with the assistance of Artinis’ fNIRS experts as part of a hands-on fNIRS workshop, with step-by-step supervision. Data were recorded using Brite Connect Ultra, an ad-hoc software platform designed for large-scale fNIRS hyperscanning, consisting of a principal recording hub and one tablet interface. This allowed two researchers (one coordinator and one assistant) to supervise the multiple simultaneous recordings. Data were recorded at a sampling frequency of 100 Hz for all participants. Event triggers were sent manually via button presses using PortaSync (a wireless remote device compatible with Artinis systems) to enable time-series synchronisation with the stimulus presentation by PsychoPy, which were delivered to a computer not integrated to the Brite system.

**Table 1.** Optode Information.

| Channel | Transmitter | Receiver | Position | Channel Length (mm) |
| --- | --- | --- | --- | --- |
| 1 | 1 | 1 | Right frontal | 25.76 |
| 2 | 1 | 2 | Right frontal | 31.62 |
| 3 | 2 | 1 | Short separation | 0 |
| 4 | 3 | 1 | Right frontal | 34.43 |
| 5 | 3 | 2 | Right frontal | 24.72 |
| 6 | 4 | 3 | Right temporal | 31.33 |
| 7 | 4 | 4 | Right temporal | 28.37 |
| 8 | 5 | 3 | Right temporal | 31.24 |
| 9 | 5 | 4 | Right temporal | 34.75 |
| 10 | 6 | 5 | Left frontal | 25.88 |
| 11 | 6 | 6 | Left frontal | 31.64 |
| 12 | 7 | 5 | Short separation | 0 |
| 13 | 8 | 5 | Left frontal | 34.46 |
| 14 | 8 | 6 | Left frontal | 24.93 |
| 15 | 9 | 7 | Left temporal | 31.39 |
| 16 | 9 | 8 | Left temporal | 28.39 |
| 17 | 10 | 7 | Left temporal | 31.15 |
| 18 | 10 | 8 | Left temporal | 34.73 |

### 2.4 Experimental procedure

The 28 audio trials were presented in a pre-determined random order, each followed by a brief break for participants, where they rated their interest in the dialogue on a scale from 1 to 5 (1 = not at all interesting; 3 = neutral; 5 = very interesting). The breaks had variable length and were controlled by the researchers to ensure that all participants had time to respond before proceeding to the next trial. On every second trial, a behavioural multiple-choice question was presented to maintain participants’ attention and engagement to the task. The question was to select descriptive keywords that best described the speech in the trial. These behavioural responses acted as measures of attention and engagement during the task but were not used in further analyses.

### 2.5 fNIRS Preprocessing and TRF Analysis

The raw fNIRS data were stored in shared near infrared spectroscopy format (SNIRF) (Tucker et al., 2022). Preprocessing was conducted in MATLAB using inspiration from the open source fNIRS preprocessing pipelines for the NIRx system^2^ and for the Artinis system using SNIRF^3^. Our pipeline requires three publicly available toolboxes: inpaint_nans (D’Errico, 2026), Homer3 (Huppert et al., 2009) and NIRS (Santosa et al., 2018). The following preprocessing steps were applied: (1) early pruning based on raw intensity levels to identify missing values and signal dropouts, (2) conversion of light intensity to optical density (OD), (3) band-pass filtering between 0.01 and 0.7 Hz was applied because stimulus-related activity is not expected below 0.01 Hz, while haemodynamic responses typically occur below 0.5 Hz. The upper cutoff also helps attenuate cardiac signals, which usually occur around 1 Hz, (4) inspection and removal of potential bad channels (none were identified), and (5) exclusion of the two short-separation channels. Subsequently, each trial was segmented and aligned with the corresponding speech stimulus, downsampled to 25 Hz, and saved in the Continuous-event Neural Data (CND) data structure (Di Liberto et al., 2024).

We extracted five stimulus features to use in our analyses including the acoustic envelope, half-way rectified envelope derivative, word onset times, and surprisal and entropy from GPT-2 (see (Ip et al., 2025) for detailed descriptions of feature extraction). Temporal response functions (TRFs) were used to estimate the linear mapping between a given sensory stimulus feature and the corresponding neural responses (Ding et al., 2014; Lalor, 2009). TRFs were computed using the mTRF-Toolbox (Crosse et al., 2016) and custom code built starting from scripts from the CNSP open science initiative^4^. TRFs estimate the relationship between the stimulus features and fNIRS signals as a linear time-invariant system via a lagged Ridge regression procedure. TRF models were fit for each participant separately, using a leave-one-out cross-validation procedure across trials to control for overfitting, and Ridge regularisation to prevent overfitting. The regularisation parameter (λ) was optimised through an exhaustive search over a logarithmic range from 0.00001 to 1000000 within each training fold. For forward models, the TRF estimates a temporal filter for each fNIRS channel, capturing how the neural response at a given time point can be predicted from the preceding stimulus. The optimal λ for forward models was defined as the value yielding the highest prediction correlation (Pearson’s r) between the predicted and observed fNIRS signals (Crosse et al., 2016). Backward models were fit to reconstruct stimulus features from the fNIRS signals, where data from all channels is considered simultaneously. The optimal value of λ for backward models was defined as the value yielding the highest Pearson’s correlation between the reconstructed and actual stimulus feature.

Since fNIRS measures haemodynamic responses evolve much slower than signals measured by EEG or MEG, several adjustments were made. First, trials were segmented by including additional 30 seconds of fNIRS data after the end of the sound stimulus, ensuring that speech-related haemodynamic responses are fully included in the trials. A technical requirement for doing so was to zero-pad the stimulus features to match the fNIRS segments. Second, we selected TRF time-lags ranging from 0 to 30 seconds to capture the slow haemodynamic response; much longer than those usually used in EEG and MEG studies. To explore the possibility of improving computational efficiency for the TRF model fit, we tested the TRF analysis for several downsampling frequencies, ranging from 1 Hz to 25 Hz, determining the minimum sampling frequency possible without information loss (Figure 1B). Finally, since fNIRS measures changes in oxygenated and deoxygenated haemoglobin concentrations (HbO and HbR respectively), separate models for fit for each metric. Due to our small sample size, all statistical comparisons were done across trials rather than across participants.

**Figure 1.**
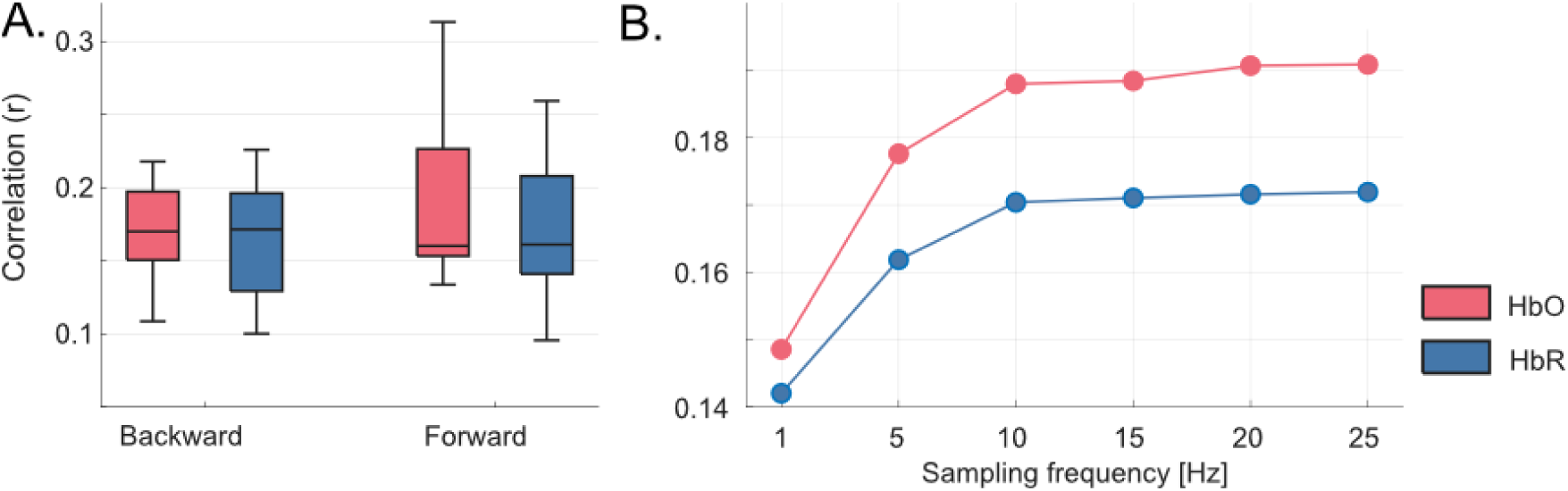
Comparison of TRF model parameters. **(A)** Backward and forward univariate TRF model performance. Red colour denotes TRF results for HbO signals, and blue denotes HbR. The y-axis refers to the Pearson’s correlation metric (envelope reconstruction and fNIRS prediction for backward and forward models respectively). **(B)** Impact of sampling frequency on the forward TRF model performance.

To test if our TRF models were explaining meaningful variance in the fNIRS response, we compared our prediction correlation values to a null distribution. For each feature and for a full multivariate model, null distributions were created by mismatching the stimulus features and fNIRS data across trials, using pairs separated by at least five trials to minimise temporal autocorrelation in the slow fNIRS signals. This was repeated 100 times per subject. Prediction correlations were computed for each iteration and averaged across channels and trials, resulting in 800 null correlation values (100 per subject). Our observed mean prediction correlations were then compared to the null distribution for the corresponding model.

### 2.6 General Linear Model analysis

To provide a benchmark for the TRF approach, we implemented a standard general linear model (GLM) analysis using a cross-validated framework. For each participant, a design matrix was constructed by convolving the extracted stimulus features (i.e., acoustic envelope, envelope derivative, word onsets, lexical surprisal, and entropy) with a canonical haemodynamic response function (HRF). The HRF was generated using standard SPM12 parameters, including a peak delay of 6 s and an undershoot delay of 16 s, and was normalised to a peak amplitude of one. Following convolution, all regressors were z-scored to normalise their scales and ensure the stability of the subsequent regression. The model also incorporated nuisance regressors to account for non-neural signal components and low-frequency fluctuations. These included trial-specific intercepts to capture baseline offsets for each speech segment and linear drift terms to model slow signal trends within each trial duration. Prior to the fitting procedure, fNIRS data were pre-processed by applying a linear detrend operation (*detrend* function in MATLAB) and z-score transformation to each individual trial.

Predictive performance was assessed using a 10-fold cross-validation procedure. At each iteration, nine segments were used to estimate the model weights via ordinary least squares (OLS) regression. The resulting weights were then applied to the design matrix of the remaining held-out segment to generate a predicted fNIRS time-series. This process was repeated for each fold, and model accuracy was quantified by calculating the Pearson’s correlation coefficient (r) between the predicted and ground-truth signals, for each of the 16 channels. These correlations were calculated separately for oxygenated (HbO), deoxygenated (HbR), and total haemoglobin (HbT) metrics to allow for a comprehensive comparison of model predictive performance across different physiological signals. By using cross-validation and Pearson’s correlations, we ensured that the GLM results were directly comparable to the TRF metrics.

## 3. Results

Participants rated the passages to be of neutral interest on average (*M* = 3.04), with the adult-child trials being rated as slightly more enjoyable (*M* = 3.20) than the adult-adult trials (*M* = 2.76). Accuracy on the comprehension questions was high (range: 76.9% - 100%, *M* = 89.4%), with accuracy being slightly higher for the adult-child trials (*M* = 91.7%) than the adult-adult trials (*M* = 87.5%).

To determine model parameters for subsequent analyses, we first explored the impact of the downsampling frequency on the performance of forward and backward models (Figure 1). We ran both backward and forward TRF models on the acoustic envelope for both HbO and HbR (Figure 1A). While backward models are often reported with higher correlation scores than forward models in EEG and MEG TRF analyses (Crosse et al., 2016; Haufe et al., 2014), we observed similar median correlations and no clear differences between modelling direction in fNIRS. Therefore, we proceeded with the forward model for subsequent analyses, primarily due to its more direct interpretability (Haufe et al., 2014). Forward models provide TRF weights that characterise how different features contribute to the fNIRS signal. Because fNIRS measures slow haemodynamic responses, lower analysis sampling frequencies than those used in EEG or MEG should be sufficient. While lower sampling rates reduce computational cost, overly low frequencies may fail to capture relevant response dynamics or, similarly, cause the loss of critical stimulus information. We tested six different analysis sampling frequencies from 1Hz to 25Hz to determine the minimum sampling frequency without information loss (Figure 1B). Performance was reduced at 1 and 5 Hz, whereas correlations appeared to stabilise at 10 Hz for both HbO and HbR. For HbO, a repeated measures ANOVA with Greenhouse-Geisser correction revealed a significant effect of sampling frequency (*F*(1.24, 33.42) = 32.22, *p* < .001). We then ran follow up pairwise t-tests (false discovery rate [FDR] corrected) on adjacent sampling frequencies (e.g., 1 to 5, 5 to 10, etc.). We found that the mean prediction correlation across trials was significant greater for 5 Hz than 1 Hz (*t*(27) = 4.77, *p* < .001) and for 10 Hz than 5 Hz (*t*(27) = 5.72, *p* < .001). There were no significant differences between 15 Hz and 10 Hz (*t*(27) = 0.35, *p* = .782, 20 Hz and 15 Hz *t*(27) = 1.63, *p* = .132, or 25 Hz and 20 Hz *t*(27) = 0.25, *p* < .808. Similarly for HbR, we found a significant effect of sampling frequency (*F*(1.32, 35.58) = 21.18, *p* < .001; Greenhouse-Geisser corrected). We then ran follow up pairwise t-tests (FDR corrected) on adjacent sampling frequencies (e.g., 1 to 5, 5 to 10, etc.). We found that the mean prediction correlation across trials was significant greater for 5 Hz than 1 Hz (*t*(27) = 3.59, *p* = .002) and for 10 Hz than 5 Hz (*t*(27) = 4.63, *p* < .001). There were no significant differences between 15 Hz and 10 Hz (*t*(27) = 0.60, *p* = .639, 20 Hz and 15 Hz *t*(27) = 0.40, *p* = .689, or 25 Hz and 20 Hz *t*(27) = 0.53, *p* < .647). Based on this trade-off between model performance and computational efficiency, subsequent analyses were conducted using a sampling frequency of 10 Hz. All subsequent results are presented for the forward model with a sampling frequency of 10 Hz, unless stated otherwise.

### 3.1 fNIRS Tracking of the Acoustic Envelope

Univariate forward TRFs relating sound envelope end fNIRS time-series were examined. HbO values yielded prediction correlations that were 10.3% higher than HbR on average (Figure 2A). Here, we also computed prediction correlations for the sum of HbO and HbR (referred to as HbT) and found comparable values to HbO (2.1% greater for HbT than HbO). As such, the analyses that follow are carried out on HbO and HbR separately. When the prediction correlations were broken down by channel, we can see that frontal channels tend to yield higher values than temporal channels for HbO (Figure 2C) but not for HbR (Figure 2D). We averaged the prediction correlation across the four channels in each group of optodes (left frontal, left temporal, right frontal, right temporal) for both HbO and HbR. For HbO, a repeated measures ANOVA with Greenhouse Geisser correction revealed a significant effect of channel location (*F*(1.76, 47.41) = 9.269, *p* < .001). Follow up paired t-tests (FDR corrected) showed that the left frontal group yielded significantly higher prediction correlations than the left temporal (*t*(27) = 3.97, *p* = .002), right frontal (*t*(27) = 2.36, *p* = .031), and right temporal groups (*t*(27) = 3.77, *p* = .002). The right frontal group led to larger prediction correlations than either temporal group (left: *t*(27) = 2.62, *p* = .029; right: *t*(27) = 2.49, *p* = .029), and there was no significant difference between the two temporal groups (*t*(27) = -0.26, *p* = .799). For HbR, a repeated measures ANOVA once again revealed a significant effect of channel location (*F*(3, 81) = 3.585, *p* = .017). Here, the only significant pairwise difference was between the left frontal and right frontal groups (*t*(27) = 3.14, *p* = .024; FDR corrected), with the prediction correlations for the left frontal channels being higher on average.

**Figure 2.**
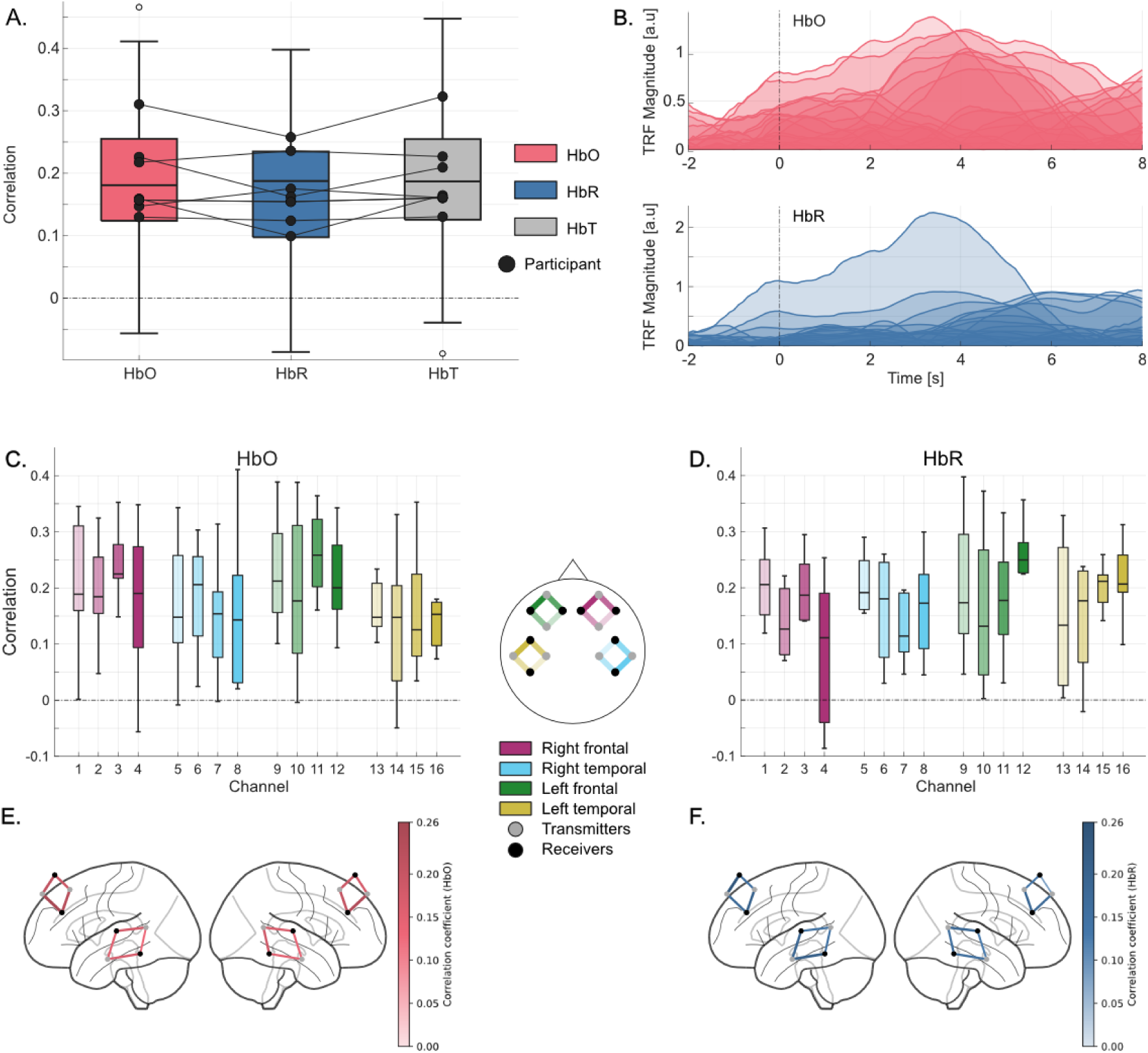
Results from forward models using the acoustic envelope speech feature at a sampling frequency of 10 Hz. **(A)** Boxplots of mean fNIRS prediction correlation values averaged across trials, channels and participants for HbO (red), HbR (blue) and HbT (grey). Each marker represents individual performance. **(B)** Magnitude of TRF weights for HbO (red) and HbR (blue). **(C-D)** Channel-wise fNIRS prediction correlation values, colour-coded by cortical region, for HbO (C) and HbR (D). **(E-F)** Channel-wise fNIRS prediction correlation values mapped onto MNI space, coloured by correlation magnitude for HbO (E) and HbR (F).

We then examined the TRF model weights for both metrics (Figure 2B). Compatibly with the properties of haemodynamic responses in the brain, which are known to peak between 3 and 6 seconds after local neural activity increases (Miezin et al., 2000), we expected TRFs to peak at latencies in that interval. Visually, the most prominent TRF components emerged at latencies of 3–5 seconds. Dominant components at those latencies emerged at seven out of 16 channels for HbO, and at one left frontal channel for HbR.

### 3.2 Univariate versus Multivariate TRFs

We next investigated how different stimulus features contribute to the TRF model. We fit five univariate models, one for each feature (envelope, envelope derivative, word onsets, lexical surprisal, and entropy), as well as a multivariate model including all five features. All TRFs led to average fNIRS prediction correlations values within 0.1 and 0.2 (Figure 3A). The univariate envelope TRF and the multivariate TRF led to the highest average prediction correlation values for both HbO (envelope: *M* = 0.188; multivariate: *M* = 0.181) and HbR (envelope: *M* = 0.170; multivariate: *M* = 0.167), while the univariate envelope derivative model resulted in the lowest prediction correlation values for both signals (HbO: *M* = 0.131; HbR: *M* = 0.122). Pairwise t-tests (FDR corrected) across all the models revealed several differences in prediction correlations. Specifically, for HbO, the multivariate model and the univariate envelope model yielded significantly higher prediction correlations than all univariate models (all *p* < .05). There was no significant difference between these two models (*t*(27) = 1.42, *p* = .167). The univariate derivative model resulted in significantly lower prediction correlations than all models except the univariate surprisal model (all *p* < .05; derivative versus surprisal: *t*(27) = -1.67, *p* = .115). For HbR, there were no significant differences between the multivariate model, the univariate envelope model, and the univariate word onset model (all *p* > .05). All three of these models yielded significantly higher prediction correlations than the univariate derivative model and the univariate surprisal model (all *p* < .05). The multivariate model and the univariate envelope model were significantly higher than the entropy model (all *p* < .05), but there was no significant difference between the univariate word onset model and the univariate entropy model (*t*(27) = 1.89, *p* = .102).

**Figure 3.**
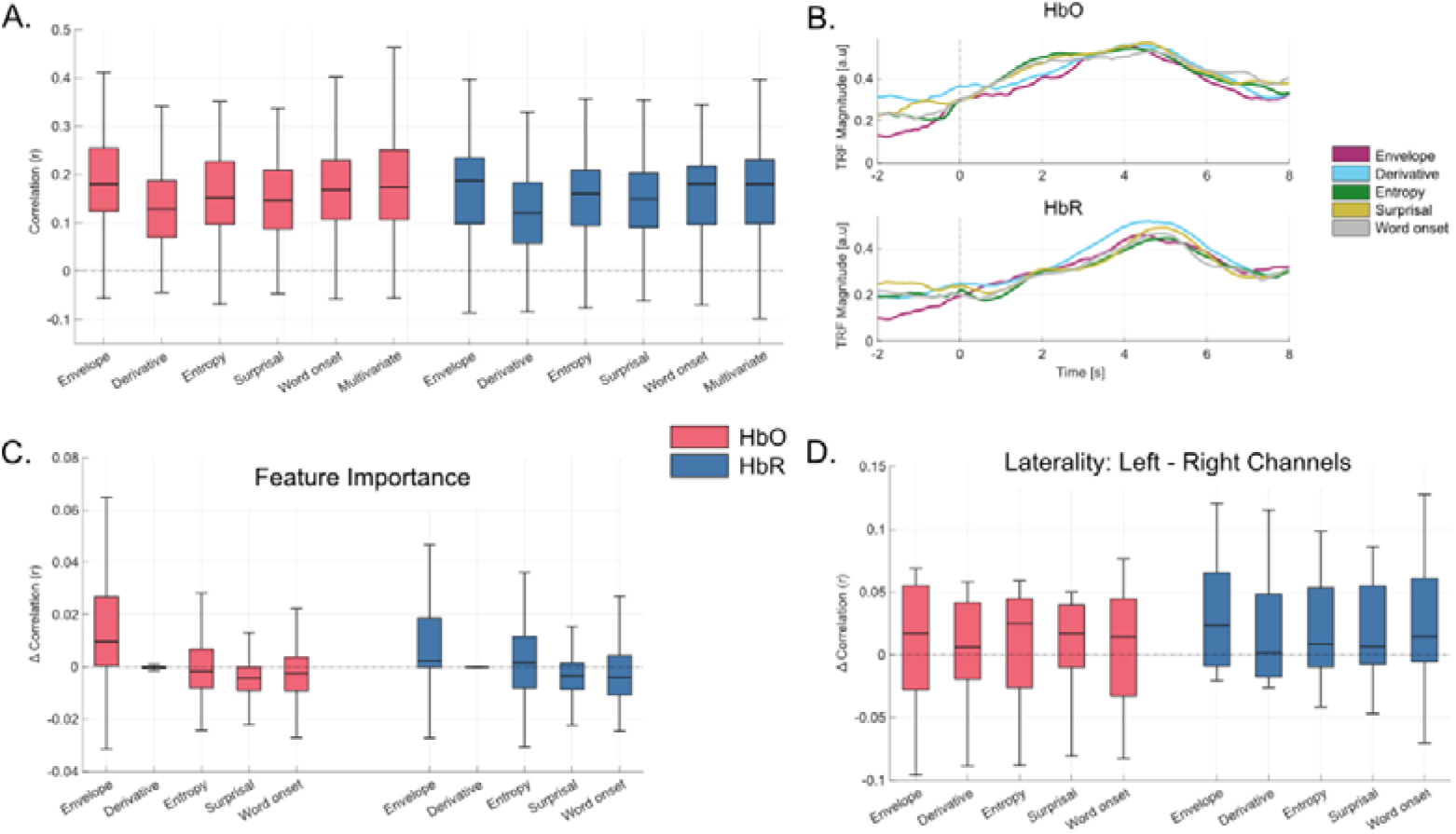
Univariate and multivariate forward TRF models at a sampling frequency of 10Hz. **(A)** Boxplots of mean prediction correlation values averaged across trials, channels, and participants for HbO (red) and HbR (blue), shown for five speech features (envelope, derivative, entropy, surprisal, word onset) as well as for a multivariate model including all features. **(B)** TRF weights averaged across trials, channels, and participants for each speech feature, shown separately for HbO (top) and HbR (bottom). **(C)** Average correlation gain of the full multivariate model (all five features) relative to multivariate models with one feature omitted. **(D)** Laterality analysis computed as the difference between fNIRS prediction correlations from left-hemisphere channels and right-hemisphere channels.

TRF weights from the multivariate model averaged across channels showed that all five speech features exhibited similar temporal profiles (Figure 3B). Based on visual inspection, for HbO the responses for all features appeared to display a broader peak between 3–5 s, whereas HbR showed a more pronounced peak around 5 s, again consistent with the haemodynamic response.

Further analyses were conducted to measure the unique contribution of each stimulus feature to the multivariate TRF (Figure 3C). That was quantified as the fNIRS prediction correlation loss after removing one stimulus feature at a time at the model training stage (i.e., feature importance). We found envelope was the largest contributor to the TRF models for both HbO and HbR. We ran paired t-tests (FDR corrected) comparing the full multivariate model to reduced models in which each feature was omitted individually. For both HbO and HbR, the only feature that resulted in significantly lower prediction correlations when dropped was the envelope (HbO: *t*(27) = 3.44, *p* = .01; HbR: *t*(27) = 2.94, *p =* .033).Together with the results from the univariate models, this shows that the envelope appears to be the most important feature in our TRF models.

The laterality of the univariate models was computed as the difference between prediction correlations from left- and right-hemisphere channels for each of speech features for both metrics (Figure 3D). We ran pairwise t-tests (FDR corrected) between the mean of the left hemisphere channels and the mean of the right hemisphere channels for each model. Although the mean difference between the left and right was positive for all features, none showed significant laterality (all *p* > .05).

### 3.4 Is it worth using TRFs to analyse fNIRS Data?

To assess whether the TRF performance exceeded chance level, we compared our obtained fNIRS prediction correlations to a null distribution of 800 datapoints created by using mismatched trial data (Figure 4A, B). TRF models resulted in significantly higher prediction correlations for all features than the null distribution (including for the multivariate model including all five features) for both HbO and HbR. For HbO, the prediction correlations we obtained from the univariate envelope, entropy, surprisal and word onset models were beyond the 99^th^ percentile of the null distribution (*p* < .01) and for the univariate derivative and multivariate model, our obtained values were beyond the 95^th^ percentile (*p* < .05). For HbR, all six models resulted in prediction correlations beyond the 99^th^ percentile of the null distribution (*p* < .01). This means that the TRF is capturing meaningful variance in response to these stimulus features for all models and both haemodynamic metrics.

**Figure 4.**
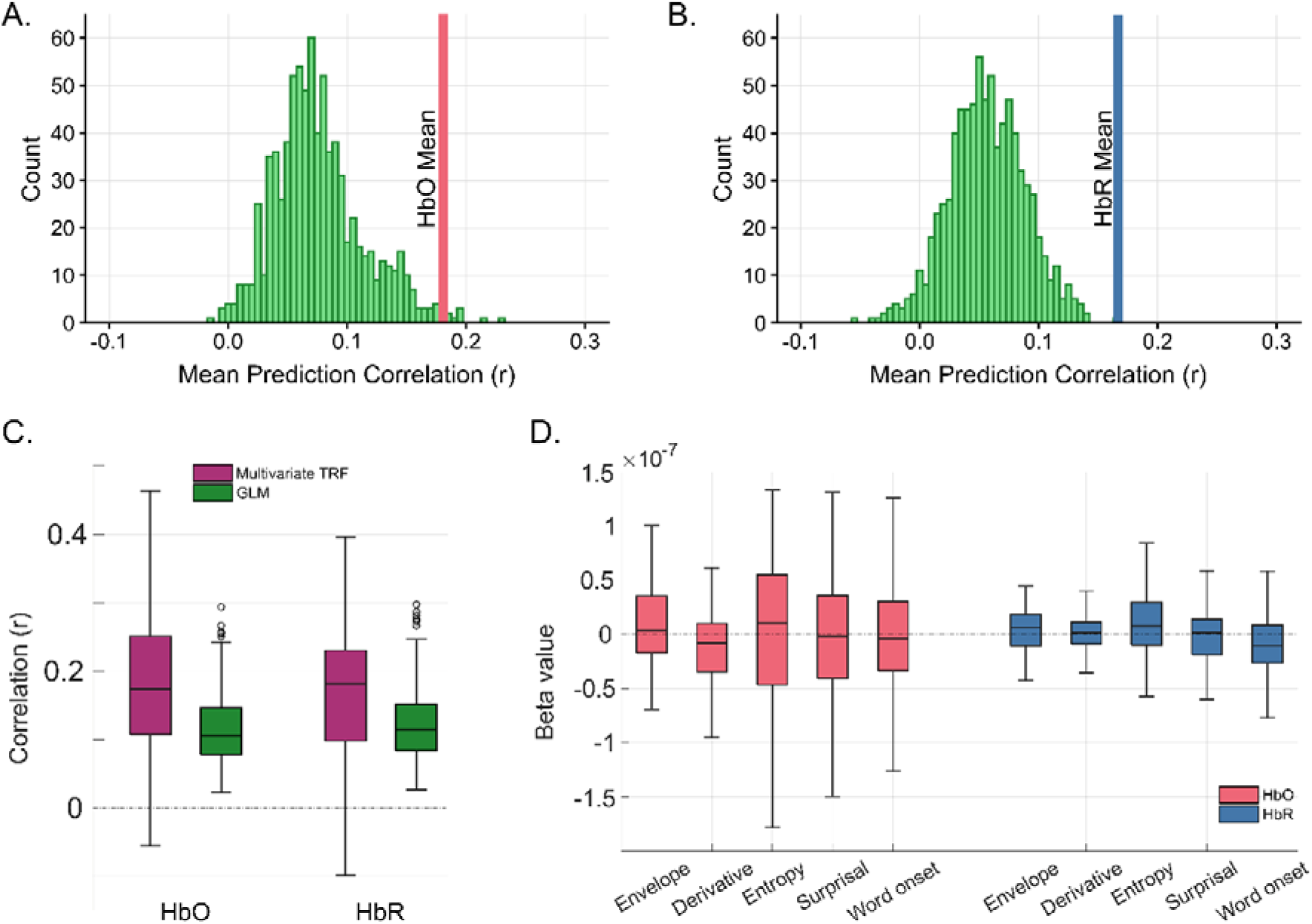
Comparison of TRF Models to Null Distributions and GLM. **(A)** Distribution of prediction correlations for the multivariate TRF model using mismatched trial data for the HbO metric. The mean prediction correlation for the actual TRF model is shown in red. **(B)** The same for the HbR metric. The mean prediction correlation for the actual TRF model is shown in blue. **(C)** Comparison of prediction correlations for the TRF (purple) and GLM (green) analyses for HbO and HbR. **(D)** Beta weights for each feature from the GLM analysis for HbO and HbR.

Next, we examined the performance of a GLM approach with cross-validation. When fitting the GLM with all five stimulus features convolved with a canonical HRF, the model successfully predicted the fNIRS signals above chance, yielding positive median correlation (r) values ranging between 0.12 and 0.13 across HbO and HbR metrics. The GLM was fitted with all five stimulus features included simultaneously. The resulting beta weights reflect the magnitude of the canonical HRF response to each feature and are conceptually comparable to the peak amplitudes of the corresponding TRFs, though unlike the TRF, the GLM does not estimate the shape of the haemodynamic response from the data. While the GLM provided a reliable linear fit, the overall predictive correlations were visibly lower than those achieved by the multivariate TRF models, which reached median correlations closer to 0.18 for both HbO and HbR. It is noteworthy that model performance was evaluated using cross-validation rather than a trial-by-trial approach; although the latter was explored, it yielded unstable predictions that did not generalize across trials, while cross-validation provides a robust and unbiased estimate of predictive performance. For this reason, statistical comparisons between the two are done at the subject level (n = 8). We used pairwise t-tests to see if there was a significant difference in prediction correlations between the multivariate TRF model and the GLM. The TRF resulted in significantly higher prediction correlations for both HbO (*t*(7) = 3.40, *p* = .011) and HbR (*t*(7) = 2.49, *p* = .041).

Analysis of the average GLM beta weights provided further insight into how individual stimulus features drove the predicted haemodynamic response within this traditional framework. Because all five regressors were included simultaneously in the model, they inherently competed to explain the variance in the continuous fNIRS signal. Within this competitive multivariate space, the acoustic envelope exhibited the most prominent positive beta weights, indicating it was the strongest unique driver of the modelled response. Conversely, the envelope derivative yielded primarily negative beta weights, and the discrete linguistic features (word onset, surprisal, and entropy) displayed weights heavily centred around zero. This suppression of non-envelope features in the GLM aligns perfectly with our TRF multivariate gain analysis. In the TRF models, acoustic envelope resulted in the highest prediction correlation, whereas adding the derivative and linguistic features yielded near-zero multivariate gain.

## 4. Discussion

The main finding of this study is that the TRF estimation method can yield meaningful and statistically significant temporal responses from fNIRS data. The estimated TRF weights present temporal dynamics that are compatible with the canonical haemodynamic response, with responses peaking at latencies between 3 and 6 seconds. These temporal profiles were consistent across stimulus features and evaluation metrics. fNIRS prediction correlations between actual and modelled signals were significantly above chance, with average Pearson’s r values between 0.1 and 0.2 across all speech features for both HbO and HbR, higher than values typically reported in EEG and comparable to those reported in MEG TRF studies. Prediction performance was also higher than that obtained using a comparable GLM approach.

A sampling frequency of 10 Hz was identified as the optimal choice for maximising computational efficiency. Instead, lower sampling frequencies led to information loss. Additionally, HbO and HbR haemoglobin metrics resulted in fNIRS prediction correlations with comparable performance, with the HbO consistently approximately 10% higher. When considering fNIRS channels separately, predictions are higher in frontal regions than temporal regions for both metrics.

### 4.1 Forward vs Backward TRF modelling

Backward (decoding) TRF models typically yield higher correlation values than forward (encoding) TRF models in neurophysiology. One key reason is that forward models evaluate the model predictions on noisy neural recordings, as the ground-truth neural signal is unknown. Backward models, instead, perform that evaluation (i.e., Pearson’s correlation) on the (clean) stimulus space, where the ground-truth is known. Another difference relates to the multivariate-to-univariate nature of the TRF mapping. Backward models consider all neural channels simultaneously, increasing the information used for the speech reconstruction compared to using a single channel. Conversely, forward models predict one channel at a time, but can combine multiple features simultaneously, which can increase the model explanatory power when features are related to complementary neural variance. With these considerations in mind, backward models are generally expected to produce prediction correlations that are higher numerically, whereas forward models would generally yield lower correlation values, but provide greater physiological interpretability (Crosse et al., 2016; Haufe et al., 2014; Wong et al., 2018).

In contrast with that literature, our results with fNIRS indicate comparable backward and forward TRF correlations (Figure 1A). One possibility is that fNIRS signals have a higher signal-to-noise (SNR), as higher SNRs increase the prediction correlation of forward models. Another possibility is that speech-related fNIRS responses are spatially localised, meaning that combining multiple channels in backward models would not increase the correlation scores. The slow characteristic of haemodynamic signals also likely plays a role in this result. In fact, such slow dynamics may be predictable using the speech envelope, while the faster envelope dynamics may be too rapid for being reconstructed with such a slow brain signal. It should also be noted that the baseline models in Figure 4A, B have mean above zero, reflecting potential similarities across trials regardless of the specific speech segment (e.g., auditory onset response). This further highlights the importance of considering a baseline model (e.g., via trial mismatching) when assessing statistical significance.

While the slow dynamics of fNIRS may be seen as a disadvantage for the reasons above, one positive implication is computational, as the time-series can be downsampled. Here, we found that the TRF analysis remains unaltered for downsampling rates down to 10 Hz, while information is lost at lower frequencies.

### 4.2 Stimulus Feature Analysis

The acoustic envelope yielded the highest fNIRS prediction correlations and contributed most strongly to the multivariate models for both HbO and HbR (Figure 3). This was expected as it is typically the case for EEG and MEG as well. An additional contributor to that strong relationship may be that the envelope is the stimulus property in our feature set with the closest rates to the fNIRS signal, possibly leading to a stronger alignment. In fact, features with fast dynamics, such as the half-way rectified envelope derivative, which have shown to be important in EEG TRF studies (Brodbeck et al., 2018; Chalas et al., 2022) had little to no contribution to the multivariate TRF.

The TRF weights did not exhibit particular variation across features (Figure 3B). The temporal dynamics for all five features resemble the canonical haemodynamic response function, with peaks between 3 and 6 seconds for both HbO and HbR. This similarity across features may explain why the multivariate gain was quite small. The implication is that, as expected from fNIRS, the spatial maps for the TRFs may be more informative than their temporal dynamics. This was seen in the univariate envelope model, with significant differences in the prediction correlations across different channels and visually different behaviour in the weights.

### 4.3 Comparison of the TRF to null distributions and GLM approaches

Adoption of an appropriate baseline model or null distribution is critical for evaluating encoding models, such as the TRF. Here, we used trial-mismatch to ensure disruption of the temporal information in the fNIRS signal. We have also explored other baselines that are common for EEG and MEG. For example, time-based shuffling methods implementing within-trial permutation or circular shifts failed to disrupt the predictive power of the TRF model, yielding surprisingly similar correlation values as the true fNIRS-stimulus trials, most likely due to the slow nature of the neural haemodynamic response measured by fNIRS. Both within-trial permutation and circular shift preserve aspects of the original time series that are statistically meaningful for the haemodynamic response, preserving the overall distribution and amplitude of the stimulus feature, with circular shift additionally preserving the autocorrelation structure. Since fNIRS signals are dominated by slow, highly autocorrelated haemodynamics, these shuffling methods can yield null model prediction correlations that are very close to those obtained with the actual stimulus features (Lancaster et al., 2018; Liégeois et al., 2021; Santosa et al., 2018). As such, we recommend considering building null distribution by combining trial-mismatch with other operations, such as time-reversal and circular shift, disrupting the stimulus-response coupling. Interestingly, the null distribution does not centre perfectly on zero. fNIRS signals at the low frequencies we included (i.e. 0.01 Hz) remain temporally smooth and strongly autocorrelated due to the sluggish haemodynamic response. Since consecutive samples are not independent and low- frequency components dominate the variance, correlation estimates from fNIRS signals can exhibit a non-zero baseline correlations (Afyouni et al., 2019; Arbabshirani et al., 2014; Olszowy et al., 2019).

Prediction correlations were higher for TRFs than the GLM analysis. This difference in predictive power reflects the increased degrees of freedom inherent in the TRF method. By modelling temporal dynamics across a range of lags, TRFs can capture complex variance in the continuous haemodynamic response that a static, canonical HRF-based GLM may underfit. Just as with our TRF analysis, the acoustic envelope was the strongest contributor to the overall model. The beta weights for all other features were near zero, with the envelope derivative resulting in negative beta weights. Together, the near-zero GLM weights for higher-level features (word onset, surprisal, and entropy) and their lack of added predictive value in the TRF approach demonstrate that the acoustic envelope dominates the trackable haemodynamic signal in this paradigm, leaving little unique variance for the other variables to explain when modelled concurrently.

### 4.4 Limitations and recommendations for fNIRS TRFs

Since we recorded data recording during the hackathon, there were several constrains. Some steps, such as mounting the optodes and placing the fNIRS cap were performed by the participants themselves, who had no previous experience with fNIRS (Yücel et al., 2025). These steps were supervised by two experts from Artinis, who gave step-by-step demonstrations and made the necessary adjustments to ensure high quality recording, which was time consuming. The environment was not as controlled as in laboratory settings. For example, external noise was minimised but could not be fully avoided. The key limitation was our sample size (N=8), which was too small for a thorough statistical testing across participants. Nonetheless, we carried out statistical tests across trials, testing for the robustness of numerical results across the different speech segments in the experiment. The limited number of participants posed limitations on certain hyperscanning-oriented analyses, such as wavelet transform coherence (WTC). That is because WTC evaluates coherence across both frequency and time dimensions, requiring long, continuous data streams to resolve slow haemodynamic fluctuations. Because our experimental trials were relatively short for targeting haemodynamic fluctuations (1-2 minutes) and featured distinct sentences, they could not be artificially concatenated or aggregated; doing so would introduce severe temporal discontinuities and edge artifacts, rendering time-resolved synchrony analyses invalid. Future hyperscanning studies aiming to use WTC should ensure continuous, long-duration stimulus presentations, larger and better selected sample providing sufficient data to improve the strength of the statistical analyses.

In our experiment, stimulus presentation and recording systems could not be directly integrated, meaning we did not have highly precise triggers. The PortaSync synchroniser is set up for triggers manually sent via a button press at the start of an event (i.e., a trial) which can creates delays. Since the fNIRS response is much slower than EEG and MEG, these delays between the actual stimulus onset and the triggers are not detrimental. Our recommendation is that the insertion of triggers for events in the recording should be automated (e.g. applying transistor-transistor logic [TTL] pulses) to ensure more consistent and better synchronisation.

We present several recommendations for future studies employing a TRF approach to fNIRS data. First, we recommend padding the stimulus and fNIRS time-series by additional 30 seconds (raw fNIRS data for the neural data and zero-padded features for the stimulus data) to allow for an entire haemodynamic cycle to take place after the trial ends. Longer time-lags than typical EEG and MEG TRFs should be considered to properly capture this haemodynamic response – up to approximately 10 seconds after feature onset. To improve computation efficiency, we recommend downsampling the neural and stimulus data down to 10 Hz. When selecting features, we recommend using those with slow fluctuations, such as the acoustic envelope over those with rapid fluctuations and sharp edges, such as the envelope derivative. Finally, when identifying a null distribution, we recommend using trial mismatch, with trials spaced far apart in time, to try to break the autocorrelation structure within the fNIRS data and minimise inflated null correlation values.

### 4.5 Conclusions

Our findings show that TRF estimation can be meaningfully applied to fNIRS data, going beyond typical application in electrophysiological recordings. Despite the slower haemodynamic signals, we were able to show that fNIRS TRFs outperform more conventional GLM based approaches. These results support the integration of continuous speech-tracking frameworks into fNIRS research and broaden the methodological tools available for naturalistic hearing and communication studies.

## CRediT Author Contributions

**Johanna Wilroth** – Data curation, formal analysis, investigation, methodology, visualisation, and writing – original draft, writing – review & editing. **Nancy Sotero Silva** – Investigation, writing – original draft, writing – review & editing. **Ali Tafakkor** – Data curation, formal analysis, investigation, methodology, visualisation, writing – original draft, writing – review & editing. **Bruno de Avo Mesquita** – Data curation, investigation, visualisation, writing – review & editing. **Emily Y. J. Ip** – Data curation, conceptualisation, methodology, writing – review & editing. **Bonnie Lau** – Investigation, methodology. **Jaimy Hannah** – Data curation, formal analysis, investigation, methodology, project administration, supervision, visualisation, writing – original draft, writing – review & editing. **Giovanni M. Di Liberto** – conceptualisation, funding acquisition, investigation, methodology, resources, supervision, writing – review & editing.

## Acknowledgements

The authors would like to thank the Cognition and Natural Sensory Processing (CNSP) initiative, for providing the blueprint for the analysis code and data standardisation guidelines used in this work. We would like to thank the organisers of the 1^st^ CNSP Hackathon and all the participants who were directly involved in the fNIRS data collection, including John O’Doherty, Cindy Zhang, and Mike Thornton. We would also like to thank Artinis Medical Systems B.V. for making this study possible, by sharing equipment, guidance on data recording in preparation for and during the CNSP-hackathon 2025.

## Footnotes

1 https://laist.com/podcasts/servant-of-pod/kids-podcasts-a-true-alternative-to-screen-time; https://creators.spotify.com/pod/profile/martine-severin/episodes/31--How-to-Become-an-Epic-ImprovComic-with-Joy-Dolo-e1js9c2

2 https://github.com/smburns47/preprocessingfNIRS

3 https://github.com/Artinis-Medical-Systems-B-V/snirf_data_example

4 https://github.com/CNSP-Workshop/CNSP-resources

